# Understanding Complex Chromatin Dynamics of Primary Human Neutrophils During PMA Induced NETosis

**DOI:** 10.1101/2024.05.31.596897

**Authors:** Brandi Atteberry, Benjamin P. Berman, Theresa K Kelly, Justin Cayford

## Abstract

**Background:** Primary human neutrophils play a pivotal role in innate immunity, mainly through the formation of neutrophil extracellular traps (NETs) in a process known as NETosis. This cell-death pathway is crucial for combating infections but is also implicated in many inflammatory diseases such as sepsis, systemic lupus erythematosus, rheumatoid arthritis, and others.

**Methods:** The study presented here investigates chromatin dynamics during NETosis by stimulating primary human neutrophils with phorbol 12-myristate 13-acetate (PMA). We adapt the ATAC-seq (Assay for Transposase-Accessible Chromatin using sequencing) method to isolated neutrophils and characterize a time-dependent chromatin response.

**Results:** We find that chromatin accessibility patterns are consistent across individual donors and most chromatin changes occur within 30 minutes, with many continuing across the 90 minutes assessed in this study. Regulatory regions gaining accessibility are associated with activity of pathways that have been implicated in NOX-dependent NET formation.

**Conclusions:** Our findings enhance the understanding of the chromatin changes underlying NETosis and also identify potential early-acting targets for modulating this process in inflammatory diseases.

## Introduction

Neutrophils are the most abundant white blood cell type in humans and play crucial roles in the innate immune system as one of the first lines of defense against infection (1). In response to pathogens, neutrophils undergo a process called NETosis which culminates in the release of Neutrophil extracellular traps (NETs) composed of chromatin fibers adorned with anti-microbial proteins and proteases that entrap and neutralize a variety of microbial invaders, including bacteria, viruses, and fungi (2-5). While NETosis represents a crucial defense mechanism, dysregulation of this process contributes to the pathogenesis of diverse inflammatory conditions, including sepsis, systemic lupus erythematosus (SLE), and rheumatoid arthritis (RA). This underscores the delicate balance required for effective immune responses, wherein the precise regulation of NETosis is vital in maintaining immune homeostasis and preventing inflammatory disease progression (6-8). Excessive NET formation in sepsis can cause tissue damage and organ dysfunction while in SLE, aberrant NETosis triggers autoimmunity by releasing autoantigens and in RA it perpetuates chronic inflammation and joint tissue destruction. Thus, maintaining the delicate balance of NETosis regulation is crucial for effective immune responses and preventing inflammatory diseases.

Understanding the mechanisms underlying NETosis is critical for identifying biomarkers and developing therapeutic interventions. However, as neutrophils are inherently designed to sense environmental perturbations, they are highly sensitive to external stimuli. Additionally, they have short half-lives and limited survival both *in-vitro* and *in-vivo*, which complicates their study (9). Their viability is further compromised by freezing, making them challenging to work with experimentally. While neutrophil-like cell lines such as differentiated HL-60’s, have been employed to investigate NETosis regulation, these models do not faithfully replicate the behavior of primary neutrophils (10). Moreover, the study of isolated neutrophils fails to capture the complexity of cell-cell interactions and indirect signaling among various immune cell types. Furthermore, findings based on studying NETosis in murine models do not always translate to human biology (11, 12). Investigating NETosis within the context of human primary neutrophils is crucial for translating laboratory-based findings into clinical advancements.

NETosis has been shown to have two distinct pathways, dependent or independent activation of NADPH Oxidase (NOX) (1, 13). Phorbol 12-myristate 13-acetate (PMA) stimulation has been shown to be NOX dependent (14), while other activators such as calcium ionophore are NOX independent (15). Both the NOX dependent and independent NETosis pathways likely contribute to host defense against pathogens, and dysregulation of either pathway can exacerbate inflammatory conditions. For instance, excessive reactive oxygen species (ROS) production in the NOX pathway can lead to tissue damage and inflammation, while aberrant activation of protein kinases in the non-oxidative pathway can contribute to autoimmune responses and tissue injury (16). Understanding the intricate regulation of these pathways is crucial for developing targeted therapeutic strategies to modulate NETosis and mitigate inflammatory diseases.

Chromatin decondensation studies have investigated the dynamics of chromatin modifications during NETosis, highlighting the importance of histone citrullination by PAD4 in chromatin decondensation and trap formation (17). Other research has explored the regulation of chromatin remodeling enzymes and their impact on NET release (18). In this study, we aimed to probe the temporal dynamics of chromatin reorganization during PMA activation of NETosis (13). Leveraging the Assay for Transposase-Accessible Chromatin using sequencing (ATAC-Seq) (19) with the incorporation of fixation methods (20), we interrogated the landscape of chromatin accessibility at distinct time points following PMA stimulation. By outlining the spatial and temporal trends in chromatin remodeling, we sought to unravel the underlying regulatory networks orchestrating NETosis and identify candidate epigenetic modifiers that may provide answers to therapeutic intervention. We found that chromatin accessibility patterns are consistent across individual donors and that chromatin dynamics were organized across the genome. Lastly, the chromatin accessibility patterns highlight key regulatory mechanisms activated during PMA stimulated NET formation.

## Materials and methods

### Tn5 Assembly

Recombinant Tn5 transposase protein (Active Motif #81284) lots were assembled with IDT custom oligos mosaic end (ME) ME_Rev, ME_A, and ME_B, and activity was tested as previously described (19) prior to initiating ATAC-seq experiments.

### Whole Blood Collection and Neutrophil Isolation

Whole blood was obtained from healthy donors in K2-EDTA tubes (BD #366643) (PrecisionMed, San Diego). Research was approved under WCG IRB Protocol number 20181025 and all human participants gave written informed consent. Each subject was self-declared healthy, between the ages of 18-50, with BMI < 30, and not taking NSAIDs. Neutrophil isolation was initiated within one hour of blood collection using the MACSxpress Whole Blood Neutrophil Isolation Kit for humans (Miltenyl Biotec #130-104-434). Briefly, a fraction of the prepared bead mixture was added to the blood sample, followed by incubation on a rotator for cell binding. Subsequent magnetic separation yielded isolated neutrophils, which were washed in 1x PBS and red blood cell (RBC) lysis to eliminate contaminants. The resulting cell pellet was resuspended in prewarmed RPMI at 37°C for induction.

### Induction with PMA/DMSO Time Course

Isolated neutrophils were resuspended in prewarmed (37°C) RPMI 1640 Medium (Gibco Cat#11875135) at a concentration of 1-2 × 10^6^ cells/mL in a final volume of 20 mL and treated with either 100 nM of phorbol 12-myristate 13-acetate (PMA) (Sigma Cat#P1585), or an equivalent volume of Dimethyl sulfoxide (DMSO) (ATCC Cat #4-X). Subsequently, each tube was incubated at 37°C with 5% CO_2_, and aliquots were collected at 30-minute intervals up to 90 minutes. Each aliquot was fixed as described below for further processing.

### Fixation

A 10x solution of formaldehyde (Formaldehyde 11%, NaCl 1M, EDTA 0.1 mM, HEPES 0.5 mM) was prepared, and a 1:10 (1% final) volume was added to the isolated neutrophils for fixation. After incubation at room temperature for 10 minutes, fixation was quenched using a 1:20 volume of glycine (2.5M), followed by incubation on ice for 5 minutes and centrifugation at 300 x *g*. Cells were subsequently washed in ice-cold 1x PBS, pelleted, flash frozen in liquid nitrogen and stored at –80°C until further processing.

### ATAC-Seq

Unfixed ATAC-Seq was performed as previously reported without major changes (21). The fixed ATAC-seq protocol was adapted from Buenrostro et al. utilizing info from Chen et al. to determine fixation steps (20, 21), where each frozen cell pellet (125,000 cells) was resuspended in 50 µl of ATAC RSB Complete (10mM Tris pH 7.5, 3 mM MgCl_2_, 10 mM NaCl, 0.1 % NP-40, 0.1% Tween-20, 0.01% digitonin) and incubated on ice for 3 minutes.1 mL of cold ATAC RSB NULL (10 mM Tris pH 7.5, 3 mM MgCl_2_, 10 mM NaCl, 0.1% Tween-20) was added, and the mixture inverted three times before pelleting nuclei at 500 x *g* for 10 minutes at 4°C. Supernatant was removed and the pellet was resuspended in Omni-ATAC Mix (22) (1xTD (10 mM Tris pH 7.5, 5mM MgCl_2_, 10% dimethylformamide), 1X PBS, 0.01% digitonin, 0.01% Tween-20) and assembled Tn5 (3.26 µM) was added to final of 50 µl and incubated at 37°C for 30 minutes in a thermal cycler and the reaction was stopped by incubating on ice for at least 5 minutes. ATAC-Seq reverse cross-link buffer (50 mM Tris pH 8.0, 200 mM NaCl, 1 mM EDTA, 1% SDS) and proteinase K (1.25 µl) (Invitrogen #25530049) was subsequently added and incubated at 65°C overnight and DNA was then isolated using Zymo DNA purification (Zymo #D4013) and eluted in pre-warmed 18.5 µl of 10 mM Tris (pH 8.0).

### DNA Library Preparation

For each sample, a DNA library was generated using NEB Next Ultra II Q5 Master Mix (NEB# M0544L) and amplified by PCR as previously described (21). PCR amplicons were purified by size selection (0.5x followed by 1.2x) using AMPure XP Reagent (Beckman Coulter #A63881), quantified using a Qubit Flex Fluorometer (ThermoFisher #Q3327) and analyzed on Agilent Tapestation 4150 with Agilent High Sensitivity D1000 tapes (Agilent #5067-5594). Each DNA library was quantified using Roche KAPA Library Quantification Kit (Roche #0796014000) and pooled for sequencing on an Illumina NextSeq2500.

### ATAC-Seq Processing and Peak Calling

ATAC-Seq alignment and processing were conducted using an ATAC-Seq Nextflow pipeline (https://nf-co.re/atacseq/2.1.2) (23) with the nf-core framework (24). Minor modifications and default settings were used unless otherwise specified. Samples were aligned to the hg38 reference genome with a fragment size parameter set to 200. Peak calling was performed using MACS2 (25) with narrow peaks identified at a false discovery rate (FDR) of 0.01 (macs_fdr=0.01).

### Untreated ATAC-Seq analysis

Consensus peaks (*.featureCounts.txt) and annotated peaks (*.annotatePeaks.txt) were generated using the nf-core pipeline and utilized for subsequent analysis. To normalize read counts across samples, scaling factors from bigwig normalization (*./bigwig/scale) (https://cran.r-project.org/web/packages/scales/index.html) were applied to balance the total number of reads across samples. For untreated samples, normalized peak counts were combined with annotated peaks to create a consensus peak dataset. UpSetR (https://cran.r-project.org/web/packages/UpSetR/index.html) plots were generated using the *.boolean.annotatePeaks.txt files. Additional metrics, including read counts, peak annotations, bigwigs, FRiP scores, insert sizes, and other alignment metrics, were generated using the nf-core ATAC-Seq pipeline.

### Treated ATAC-Seq analysis

Treated samples were analyzed similarly to untreated samples with some modifications. Samples were normalized in the same manner as above and then were divided into six groups based on the time course (T30, T60, and T90 minutes) and treatment (DMSO vs PMA). Using *.boolean.annotatePeaks.txt files, consensus peaks were retained if at least 2/3 of the donors had a peak in that interval. The consensus peaks were then concatenated, retaining only the unique intervals. The normalized counts for these consensus peaks were then analyzed using DESeq2 (26). The DESeqDataSetFromMatrix() function was used to create the dataset, followed by the assay() function for data extraction. Principal Component Analysis (PCA) scores were calculated using prcomp, and PCA contributions were calculated with the summary() command. Pairwise comparisons were performed with DESeq(), and results were summarized using the results() function. Significant regions (padj < 0.01) were z-score normalized based on the normalized peak counts. Heatmaps were generated using pheatmap (https://cran.r-project.org/web/packages/pheatmap/index.html) with row clustering. Volcano plots were created from the results() function from DESeq2, with significant regions defined by -log_10_(padj) > 20. For the all-vs-all comparisons, significant regions from each grouping (treatment and timepoint) with padj < 0.01 were included. Pheatmap was then used with both row and column clustering. To analyze overlaps between DMSO Early vs PMA Early, Mid, and Late, an inner_join (tidyverse) (27) was used, followed by eulerr version 7.0.2 (https://CRAN.R-project.org/package=eulerr) for venn diagram visualization in a custom R script.

### HOMER Analysis

To determine if there was any enrichment of motifs in the gained or lost peaks after PMA stimulation, HOMER(28) analysis was completed on the top 1,000 significant regions based on Wald scoring for all PMA or DMSO treatments (Early, Mid, and Late). The findMotifsGenome.pl with hg38 with standard settings were used.

## Results

### Chromatin structure of primary human neutrophils is organized and stable with common accessibility patterns across healthy donors

To gain a deeper understanding of the chromatin structure of neutrophils, we performed ATAC-Seq with formaldehyde fixation of primary human neutrophils (Figure 1A) in 6 healthy donors and standard (un-fixed) ATAC-Seq in 1 healthy donor (Figure 1B). To assess the open chromatin regions across donors, we looked at genes associated with the activating H3K4me3 modification across various stages of neutrophil development (29) (Figure 1B). We found the expected overlap between identified H3K4me3 regions and ATAC-Seq peaks in our samples and what has been reported for polymorphonuclear neutrophils (PMN) (30) indicating that there is stable organization of accessible chromatin in neutrophils, across donors. Fixed ATAC-Seq samples from donors (D35, D39, D47, D58, D71, and D73) had higher signal overall compared to the unfixed sample (D32) as is highlighted in the *CLEC7A* and *HCAR* loci (Figure 1B). There was also signal in those loci in D32 but the signal to noise was much lower. D32 also had signal present in regions such as *AZU1, MPO*, and *CTSG* which are more associated with promyelocytes and are not represented in the fixed samples (Figure 1B).

**Figure 1.**
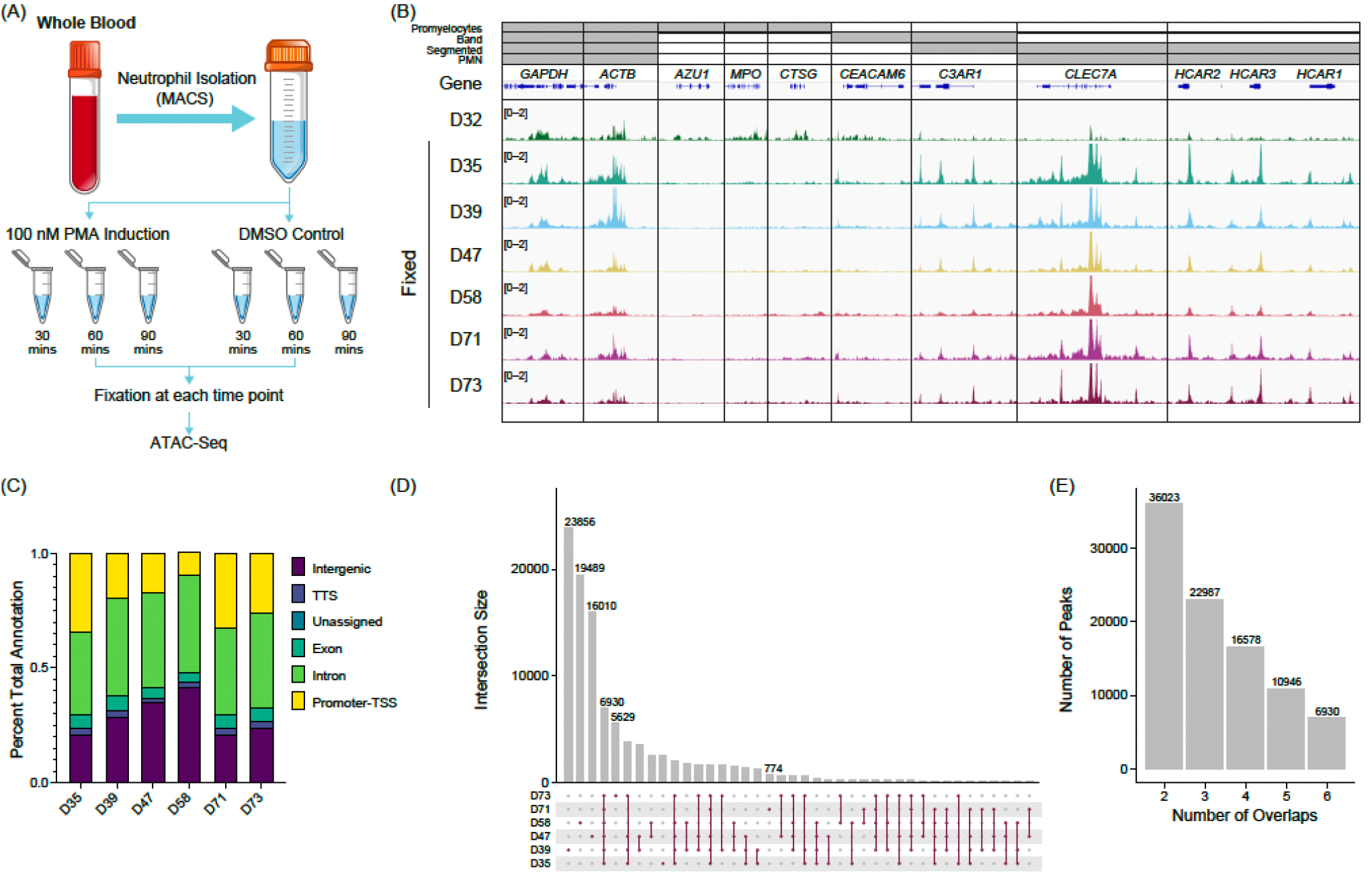
Untreated primary human neutrophil ATAC-Seq shows consistent peaks: **(A)** Experimental schematic. Fresh whole blood was collected, and neutrophils were isolated using the MACSxpress Whole Blood Neutrophil Isolation Kit for humans (Miltenyi Biotec #130-104-434). Neutrophils were then stimulated with phorbol 12-myristate 13-acetate (PMA) or DMSO for a given time, formaldehyde-fixed, and followed by ATAC-Seq. **(B)** Merged replicate IGV tracks with untreated healthy donors (n=7), standard ATAC-Seq (D32 – top), and fixed samples (D35, D39, D47, D58, D71, and D72) are shown below. The top bars (grey) indicate the stage of neutrophil development where H3K4me3 signal was present (promyelocytes, band neutrophils (band), segmented, or polymorphonuclear neutrophils (PMNs)). Housekeeping genes (*GAPDH* and *ACTB*) are shown, followed by various other neutrophil genes (*AZU1, MPO, CTSG, CEACAM6, C3AR1, CLEC7A*, and *HCAR1, HCAR2*, and *HCAR3*). **(C)** Peak annotations generated on called MACS2 peaks (q < 0.01) for the fixed donors shown as a percentage. **(D)** UpSetR plot for the fixed donors showing the intersections of untreated neutrophils. **(E)** Total number of consensus peaks (y-axis) based on the overlap for the indicated number of donors (x-axis) for the fixed samples (n=6).

We identified peaks using MACS2 (q < 0.01) and found that accessibility regions were present across a variety of genomic features with some variability across samples (Figure 1C), which potentially was driven by donor-to-donor differences. We found the peaks were reproducible across the 6 healthy donors indicated by a high number of consensus peaks. Donors D39, D47, and D58 had a high number of unique peaks (Figure 1D), which may be due to variable sequencing depth among samples (Supplementary Figure 1A). There is an overall trend of increased FRiP as the number of peaks increase with some variability, which is not unusual when using primary cells (Supplementary Figure 1B). The number of peaks shared across all six fixed donors was 6,930 (Figure 1E). The unfixed sample, D32, had significantly more peaks than the fixed samples (>100,000) and most of the peaks did not correlate with the fixed samples. Of the approximate 7,000 peaks found in all fixed samples, over 81% were also shared with the unfixed donor (Supplementary Figure 1C). To generate a consensus neutrophil peak set across the donors, we set the threshold of the peak being present in at least half of the fixed samples for the peak to be included as a consensus peak. We found around 23,000 peaks were present using this threshold (Figure 1E). These findings suggest a well-structured and regulated chromatin landscape of primary human neutrophils.

### Chromatin dynamics occur throughout PMA stimulation compared to DMSO controls

To assess chromatin changes associated with PMA induced NETosis we treated isolated neutrophils from 3 donors with PMA or DMSO and collected samples at 30, 60, and 90 minutes. There was a reproducible response to PMA stimulation but not with a DMSO vehicle control compared to the untreated samples (Figure 2A). We found consistent chromatin accessibility in house-keeping genes: *TBP, RPL32* (Figure 2A), *GAPDH*, and *ACTB* (data not shown) throughout the untreated, DMSO, and PMA treatments. Interestingly, peaks were both gained or lost upon PMA stimulation. A gain of peaks generally appeared within 30 minutes (Early) of stimulation as highlighted at the *RAD9a* and *KCANB2* loci (Figure 2A). Loss of peaks started to appear at the Early time-point but were not fully resolved until 60-90 minutes (Mid-Late). The *HCK* and *TNFAIP6* loci are two examples which showed this reduction of ATAC-Seq signal over the time-course reflective the general genome-wide trend (Figure 2A).

**Figure 2.**
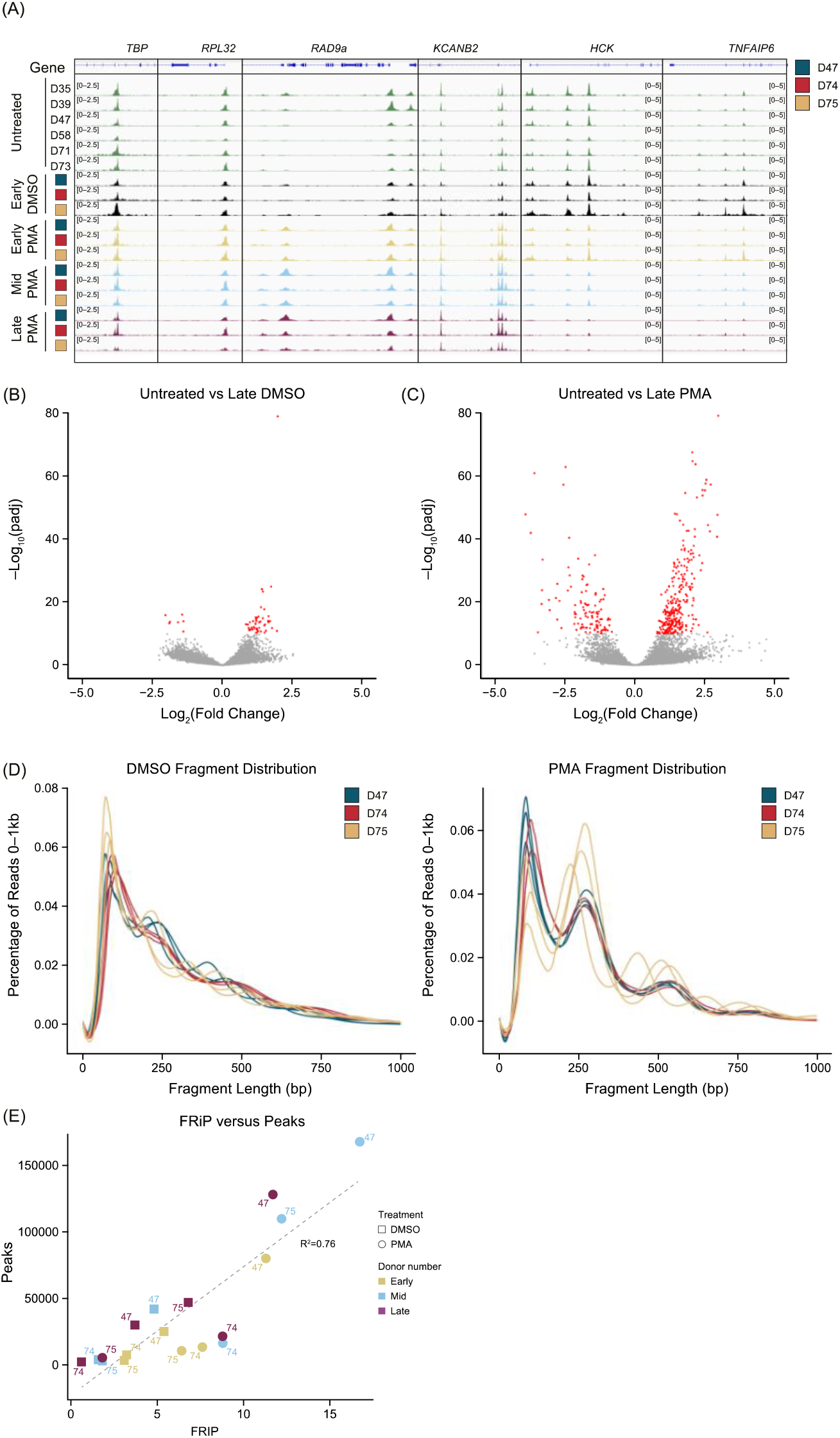
PMA generates unique chromatin dynamics compared to DMSO or untreated neutrophils: **(A)** Merged replicate IGV tracks with untreated healthy donors (n=6, green tracks) and PMA or DMSO treated donors (n=3) at Early (T30 minutes – yellow tracks), Mid (T60 minutes – blue tracks), and Late (T90 minutes – purple tracks). The treated donors D47 (blue), D74 (red), and D75 (gold) are shown at known housekeeping genes: *TBP* and *RPL32*. Examples of gained peaks are shown at the *RAD9a* and *KCANB2* loci. Decreased peak examples are shown at the *HCK* and *TNFAIP6* loci. **(B)** Volcano plot comparing untreated samples with Late DMSO. DESeq2 was used for a pairwise comparison and then plotted. The -log_10_(p.adj) value is graphed on the y-axis and significant values are plotted in red (-log_10_(p.adj) < 10). Log_2_(fold change) is indicated on the x-axis. **(C)** Similar to Figure 2B but comparing untreated samples vs. Late PMA. **(D)** Sequencing fragment distribution for the DMSO controls (left) and PMA samples (right) plotted as base pairs (bp) on the x-axis from 0-1,000, grouped into 20 bp windows. The percentage of total reads over 0-1,000 is plotted on the y-axis. Donors are indicated by color (D47 – blue, D74 – red, and D75 – gold). Timepoints (Early, Mid, and Late) are graphed per donor. **(E)** A comparison of FRiP (x-axis) with the total number of MACS2 peaks (q < 0.01) for each sample (merged replicates). DMSO samples are indicated by squares and PMA samples are indicated by circles. The timepoint is represented by color (Early – gold, Mid – blue, Late – purple). Each donor number is indicated, and the dashed line represents y=x. Correlation (R^2^) was 0.76.

We found that DMSO treated samples had a low number of differential accessible regions (DARs) at the most extreme pairwise comparison (DESeq2) of untreated versus Late DMSO (90-minute incubation) (Figure 2B). However, there were many DARs between the untreated the Late PMA treatment across the 3 donors (Figure 2C). Volcano plots, show a much greater response overall in the PMA comparison in both the p-values present and the fold change. As a result, we used Early DMSO treated samples as the analysis control.

Interestingly, a shift towards mono- and di-nucleosome fragments in the DNA fragment sizes was observed after PMA treatment at all time-points (Figure 2D), indicating a dynamic chromatin response. Despite the nucleosome pattern, PMA stimulation did not have an impact on peak localization (Supplementary Figure 2B), as there was a larger impact of donor compared to stimulation condition or time-point effect.

The Early PMA conditions had an increased number of peaks compared to the DMSO control at the same time. We found donors D47 and D75 had a large increase in the number of peaks following PMA treatment from Early to Mid time points, followed by a reduction from Mid to Late (Figure 2E). This suggests the loss of peaks happened at a later stage of NETosis and the increased accessibility occurred more rapidly, within the Early to Mid-timepoints. Interestingly we observed a disproportionate increase in FRiP in D74 and D75 (Figure 2E).

### PMA induction of NETosis leads to an organized chromatin response across time and donors

To determine if PMA stimulation responses were unique compared to DMSO, we performed unsupervised clustering Principal Component Analysis (PCA) comparing PMA to DMSO using unmerged replicates (Figure 3A). The first principal component (PC1) accounted for 65% of the proportion of variance between PMA treatment and DMSO vehicle controls and PC2 accounted for 10.4% of the variance and separated each group based on time (Early, Mid, and Late). Importantly, we did not observe any obvious donor batch effects (Figure 3A).

**Figure 3.**
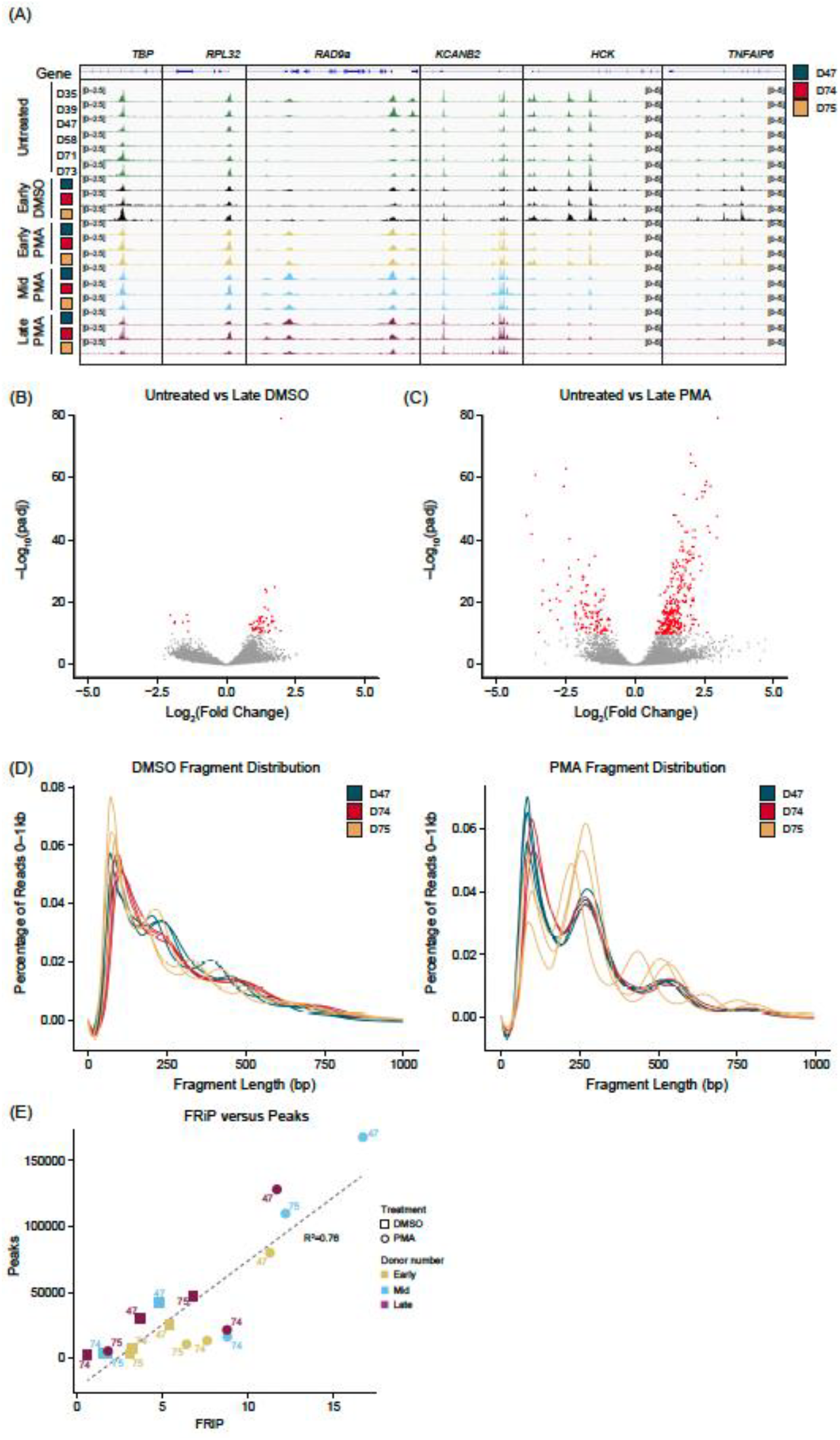
PMA stimulation causes a genome-wide alteration of chromatin accessibility across donors and time: **(A)** Principal Component Analysis (PCA) based on unbiased clustering of DMSO versus PMA using the unmerged data. Timepoints are indicated by color (Early – gold, Mid – blue, and Late – purple). Treatment is indicated by squares (DMSO) or circles (PMA). PC1 (x-axis) represented 65% of the total variance and PC2 (y-axis) represented 10.4% of variance. Donors are indicated by the number adjacent to the datapoint. **(B)** Merged replicate IGV tracks with Early DMSO (black), Early PMA (gold), Mid PMA (blue), and Late PMA (purple) at various gene loci. Donors are indicated by color (D47 – blue, D74 – red, and D75 – gold). Significant differential accessible regions based on Early DMSO vs. each time point (DESeq2 p.adj > 0.01 and log2(fold change) less than −1.5 or greater than 1.5) are shown below the genes, where gained peaks are shown in red and loss of peaks are shown in blue. Bimodal response is shown at *ACTG1* and *RAB7A*, upregulation in *PRKCB, RPLP2*, and *PNPLA2*, and downregulation in *CYTIP, RGS2*, and *ALOX5*. **(C)** Heatmap showing the number of differential accessible regions (DARs) between all pairwise comparisons (each timepoint for each treatment condition) (DESeq2 p.adj > 0.01 and log2(fold change) less than −1.5 or greater than 1.5). **(D)** Venn Diagram of the overlaps between Early DMSO and the PMA treatments (Early – gold, Mid – blue, Late – purple). The total number of DARs in each condition is indicated. **(E)** Z-score normalized heatmap representation of the Early DMSO vs. PMA treatments (Early – left, Mid – middle, Late – right). Donors are indicated by color (D47 – blue, D74 – red, and D75 – gold). Timepoints (Early, Mid, and Late) are graphed per donor. Treatment is indicated by DMSO (blue) and PMA (red). The top 1,000 DARs were selected based on the Wald score. Columns are organized by treatment, timepoint, and donor across all conditions, and rows had hierarchical clustering.

There are several examples of chromatin accessibility changes in inflammatory genes or genes known to be important in neutrophil functions (Figure 3B), with a few different modes of chromatin accessible changes observed. Regions, such as *ACTG1* and *RAB7a*, were more accessible in the Early and into the Mid stage followed by less accessible at the Late timepoint. We found some instances of intragenic upregulation (*PRKCB*) or at transcription start sites (*RPLP2* and *PNPLA2*). Also, there were regions which had a reduction in peaks (*CYTIP, RGS2*, and *ALOX5*) (Figure 3B). These represent the dynamic and structured chromatin response seen genome-wide during PMA activation across donors.

We next performed pairwise comparisons (DESeq2) between every grouping (each timepoint and treatment) and calculated the total number of Differentially Accessible Regions (DARs) (q < 0.01). The DARs highlighted large differences between the PMA and DMSO groupings but very minor differences within treatment groups (Figure 3C). Due to limited variability in the DMSO treatment group, future comparisons utilized only the Early DMSO as the control. Overlapping DARs between the PMA groups (Early, Mid, Late) were identified using a q-value < 0.01 and a log_2_ fold change greater than 1.5 or less than −1.5 (Figure 3D). Almost 40% (1121/2922) of the DARs were sustained in at least two of the time points, with the majority of regions being found in the Early or Mid-timepoints. These data suggest an initial response at 30 minutes, followed by a secondary response at 60 minutes. After 60 minutes, there were less than 5% (117/2922) unique changes (Figure 3D). Next pairwise comparisons of the Early DMSO vs each PMA treatment time and were compared. The top 1,000 DARs were z-score normalized and plotted as a heatmap (Figure 3E). The results suggest the stability of the DARs over time and across donors over each comparison.

### PMA induction has unique Early and Mid-chromatin responses associated with NETosis TFs with limited changes after 60 minutes

To obtain the highest significant DARs based on their Wald test score, the top 15% (500) upregulated in early DMSO and the top 500 DARs for all PMA timepoints were selected. The DARs were then z-score normalized and ordered based on treatment and time (Figure 4A). There was some variability in the PMA DARs with the largest differences being between Early PMA and Mid PMA. The data was further reduced to the top 1.5% DARs (50) in DMSO or PMA to highlight the largest differences between the treatments and timepoints. There are numerous DARs which appear Early and then are lost as well as the opposite of a lack of signal increasing into the Mid timepoint (Figure 4A – right). These data also support the similarities between the Mid and Late PMA treatments.

**Figure 4.**
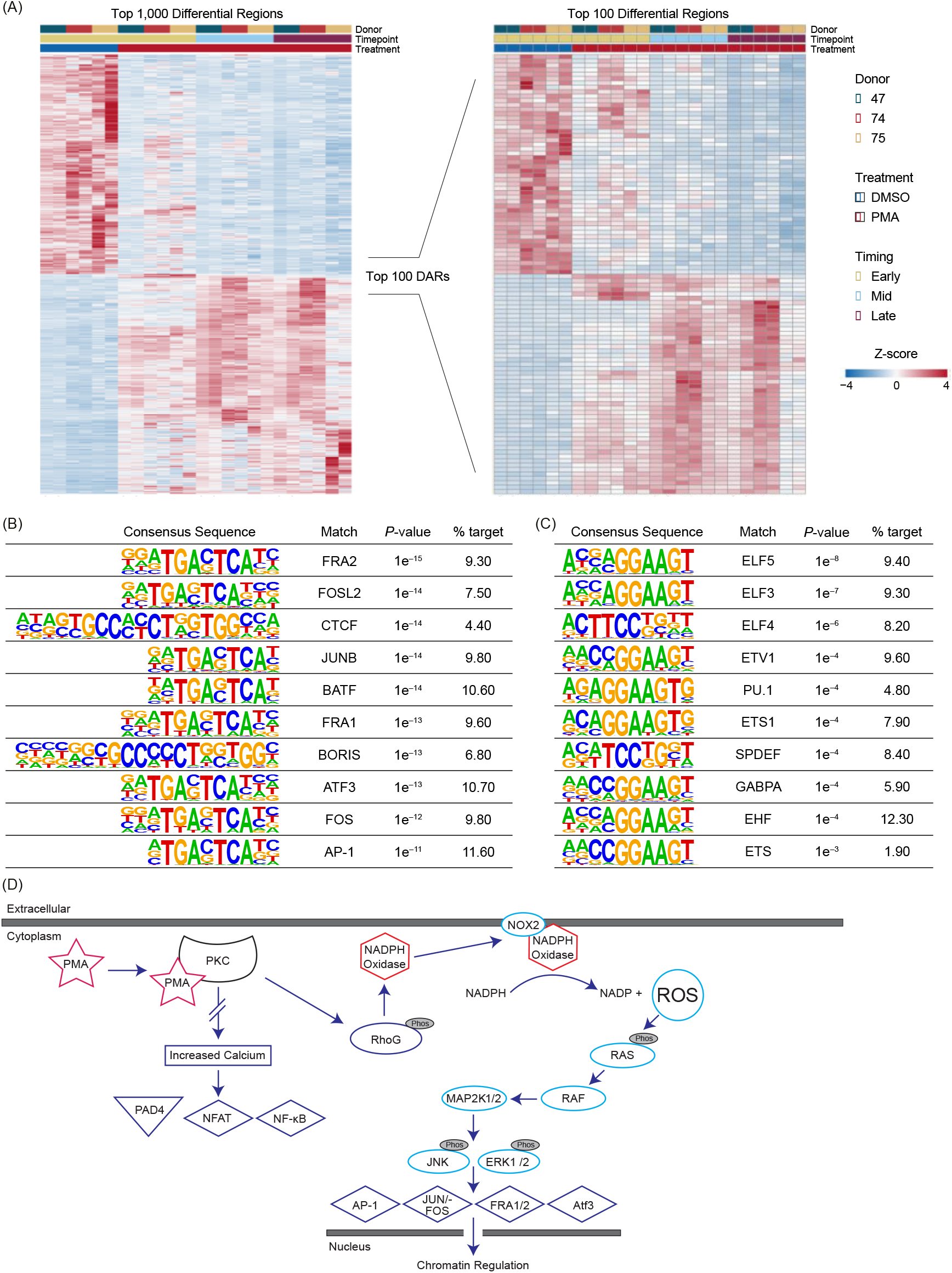
PMA-induced NETosis leads to upregulation of chromatin accessibility in RAS-associated transcription factors: **(A)** Z-score normalized heatmap representation of the top 500 (left panel) differential accessible regions (DARs) for DMSO Early and PMA based on Wald score (DESeq2 p.adj > 0.01 and log_2_(fold change) less than −1.5 or greater than 1.5). The top 100 DARs (right panel) were then filtered by Wald score (50 upregulated in DMSO and 50 upregulated in PMA). Donors are indicated by color (D47 – blue, D74 – red, and D75 – gold). Timepoints (Early, Mid, and Late) are graphed per donor. Treatment is indicated by DMSO (blue) and PMA (red). Columns are organized by treatment, timepoint, and then donor across all conditions, and rows had hierarchical clustering. **(B)** HOMER analysis from the top 1,000 upregulated DARs in PMA samples (DESeq2 p.adj > 0.01 and log2(fold change) > 1.5). **(C)** Similar to Figure 4B but upregulated in DMSO samples (log_2_(fold change) < 1.5). **(D)** Potential mechanism for transcription factor (TF) activation through PMA-activated NETosis. Briefly, PMA activates PKC, which causes activation of NADPH and NOX through RhoG to generate an increase in intracellular reactive oxygen species (ROS). ROS then activate the RAS/RAF pathway and lead to downstream activation of TFs such as AP-1, JUN/FOS, FRA1, FRA2, and ATF3. There is no increased calcium level caused by PKC activation, so PAD4, NFAT, and NF-κB are not activated.

The top 1,000 DARs from DMSO or PMA were then used for HOMER analysis (Figure 4B and Figure 4C). In the PMA DARs, transcription factors (TFs) previously associated with NETosis and inflammation, such as Fra1 (31, 32), Fra2 (13), Jun/Fos (13), Atf3 (33), and Ap-1 (13) were among the more prevalent (Figure 4B and Figure 4D). CTCF was also present in 4.4% of target sequences of the PMA DARs, which indicated a potential role of CTCF in the regulation of the chromatin response, however CTCF is often overrepresented in HOMER analysis (34) so future studies should probe the potential impact of CTCF in chromatin regulation during NETosis. In DMSO DARs, there was not a strong correlation to Netosis associated TF binding sites or many direct pathways (Figure 4C). However, one of the top HOMER results was from the ELF family, which has been associated with autoimmune disease and reduction of ELF4 in macrophages has been correlated to increased neutrophilic infiltration (35, 36).

Overall, TF binding sites associated with PAD4 such as NF-AT and NF-κb were not well represented in up or down regulated DARs (Figure 4B and Figure 4C), which could be modulated by a lack of intracellular calcium or other mechanistic reasons (Figure 4D). The representation of RAS associated TFs seen in Figure 4B support previous findings of RAS activation through PMA stimulation (13, 37). These results highlight the importance of these TFs and their potential roles in NETosis chromatin regulation (35).

## Discussion

Deciphering the molecular choreography underlying NETosis will provide important insights into the mechanisms governing both host defense and inflammation-associated pathology. Changes in chromatin architecture are an important early step in altering gene expression and initiating cellular regulatory programs, likely dictating the fate of neutrophils and shaping the ensuing immune response.

We developed an ATAC-seq approach for isolated primary human neutrophils and found that accessible chromatin regions were consistent across multiple donors, similar to other primary immune cell types (22, 38). We observed some donor-to-donor variability, but we generated a set of over 22,000 consensus peaks that were shared among 50% of the donors tested.

PMA is a strong inducer of NETosis through activation of protein kinase C (PKC), which leads to increased intracellular reactive oxygen species (ROS) through NADPH oxidase (NOX) (13, 14). We used our ATAC-seq method to study PMA-induced neutrophils, first showing that DMSO-treated neutrophils were similar to untreated samples for up to 90 minutes post-treatment (Figure 2B). When neutrophils were treated with PMA, there was a consistent response across donors at multiple timepoints, suggesting an ordered and mechanistic regulatory program. A unique change in fragment profile could be seen as early as 30 minutes (Figure 2D) suggesting that chromatin re-organization happens prior to NET production which can take several hours following PMA stimulation (39).

In concert with this early fragmentation change, we identified about 1,500 differentially accessible regions (DARs) that occurred between PMA and DMSO treated cells at 30 minutes, and the majority of these remained so up to 90 minutes. Globally, PMA-treated samples clustered away from DMSO-treated samples in an unsupervised clustering analysis (PCA), with sub-clustering by time point among the PMA-treated samples and limited donor effects. Protein Kinase C -β (*PRKCB*) has previously been shown to be upregulated during PMA stimulation (14, 40) and showed a stable increase in chromatin accessibility. More complex chromatin accessibility changes across the time course were evident at loci such as *ACTG1*, which has been shown to be silenced through the MAP2K/ERK signaling pathway (41), a pathway that is upregulated upon PMA stimulation of neutrophils (42).

Interestingly we observed a gain of accessibility at DARs containing binding sites for the transcription factors JUN/FOS, FRA1, FRA2, ATF3 and AP-1 (13, 31-33, 40, 43, 44) (Figure 4B). Upon PMA stimulation, an increase of ROS intracellularly can lead to various effects including the activation of the RAS pathway and activation of these same TFs, but not other TFs such as PAD4, NF-AT, or NF-kB (13, 14, 31, 32, 45). There was not a strong signal for TFs associated with reduced chromatin accessibility, with the most significant hits occurring in the ELF TF family, which are known to be involved with many processes including cell proliferation (46). A potential reason for a weaker signal in TFs associated with the loss of accessibility could be that the decondensation of the chromatin is targeted through a variety of TFs or another mechanism. The transcription factor motif findings support previous data that PMA stimulated neutrophils act via a NOX dependent mechanism, which activates the RAS pathway (Figure 4D) as opposed to when NETosis is activated through the NOX independent pathway, which leads to activation of PAD4 and subsequently NFAT and NF-kB (7, 14, 40). Importantly, we did not identify NFAT or NF-kB binding sites in our HOMER analysis of the top 1,000 DARs (Figure 4B-C).

Our findings not only deepen our comprehension of the epigenetic blueprint governing PMA stimulated NETosis, but also hold promise for the development of targeted strategies to modulate the earliest stages of this process in the context of inflammatory diseases.

## Supplementary

**Supplementary Figure 1:**
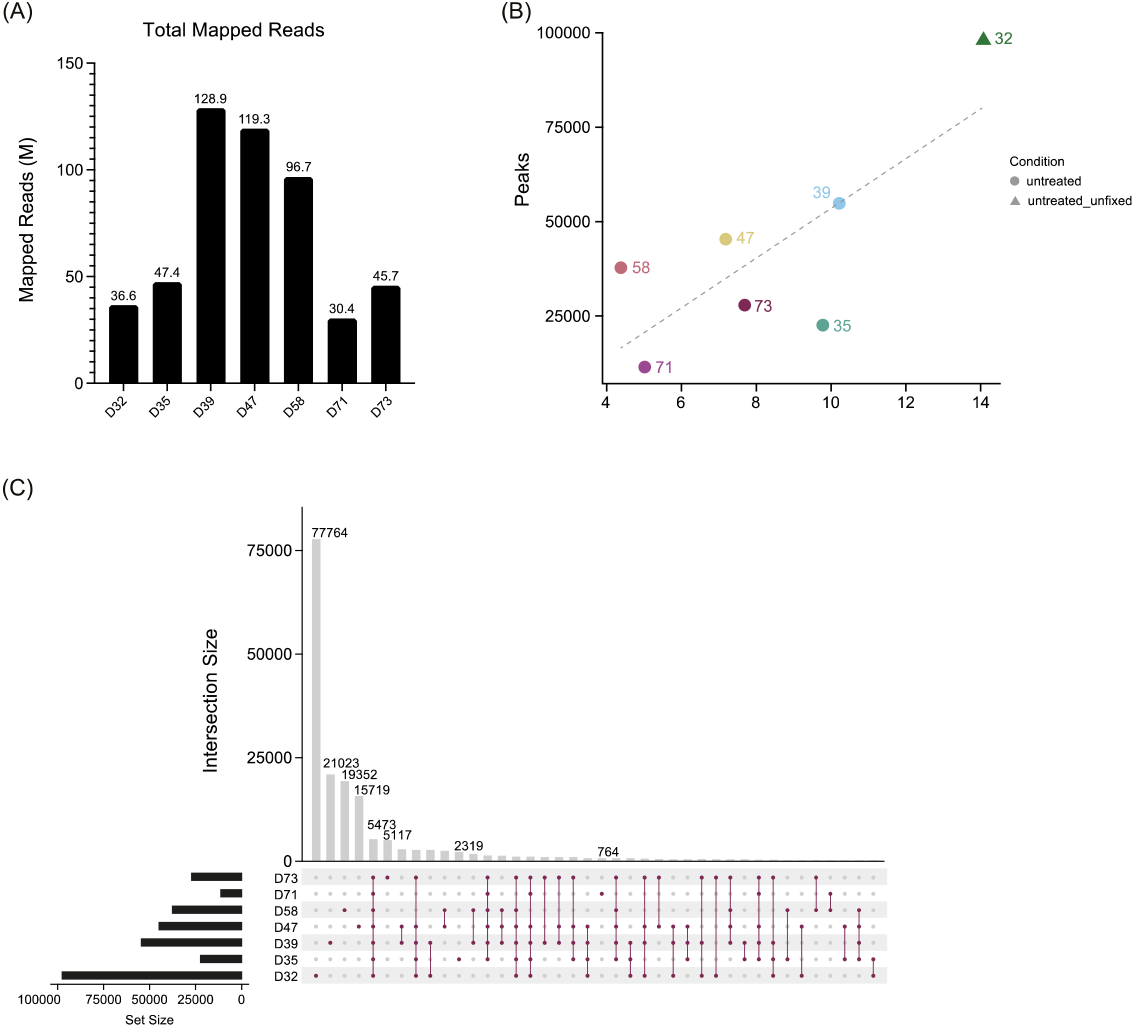
**(A)** Total number of merged mapped reads for the samples. The donors are indicated on the x-axis, and the number of mapped reads (in millions) is on the y-axis. **(B)** A comparison of FRiP (x-axis) with the total number of MACS2 peaks (q < 0.01) for each sample (merged replicates) including D32. Fixed samples are indicated by circles, and the unfixed sample is indicated by a triangle. Each donor number is indicated, and the dashed line represents y=x. Correlation (R^2^) was 0.60. **(C)** Similar to Figure 1D but D32 (unfixed sample) was included.

**Supplementary Figure 2:**
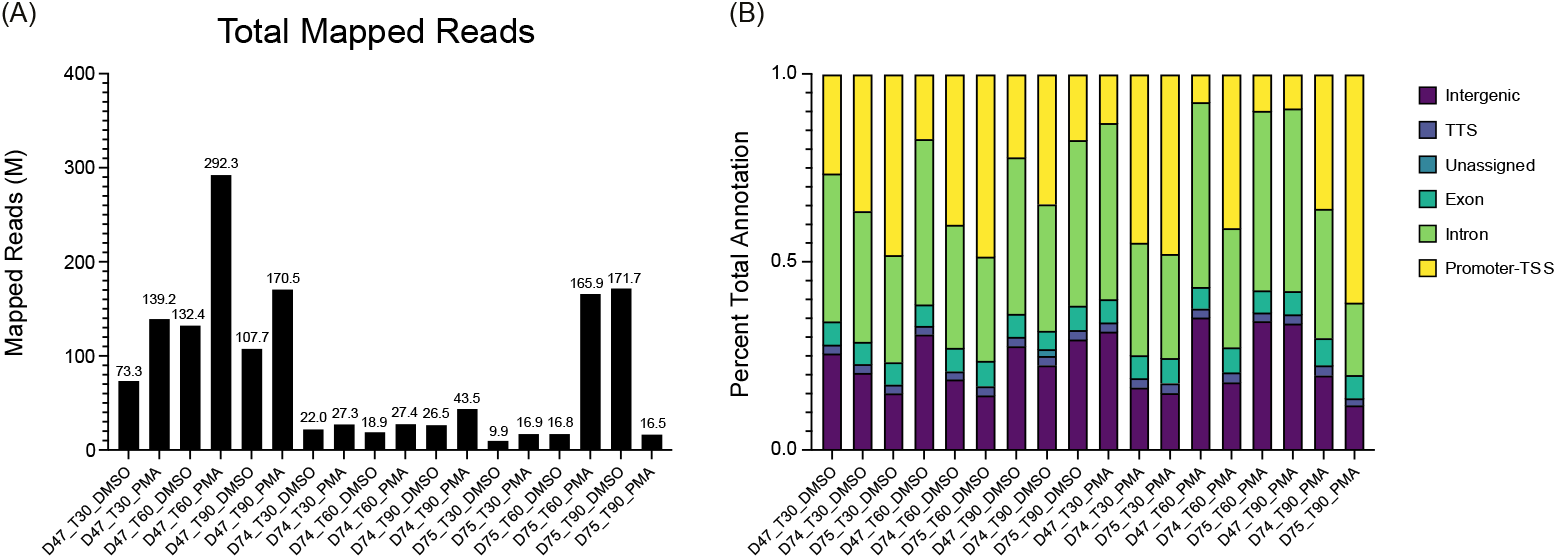
**(A)** Similar to Supplementary Figure 1A but using the merged treated samples. **(B)** Similar to Figure 1C but using the merged treated samples.

## Tables

### Supplementary Tables.xlsx

Tables including sequencing statistics, donor information, and the top 1,000 differentially accessible regions (DARs) based on the Wald test (stat column)

## Conflict of Interest

Conflict of Interest Disclosures: All authors are employees or contractors for VolitionRx. JC, BPB, TKK, and BA hold stock in VolitionRx.

## Author Contributions

JC, TKK and BPB conceived and designed the study, BA performed the experiments, JC provided guidance and JC and BPB performed data interpretation. All authors participated in interpretation of data and critical revision of the manuscript for important intellectual content.

## Funding

This work was funded by VolitionRx.

## Acknowledgments

We would like to thank Sarah Erdman, Finley Serneo, Akanksha Singh-Taylor and Kieran Zukas, as well as the broader scientific team at Volition for helpful review of the data and discussion. ChatGPT version 4o was used to support the writing of R codes and minor edits throughout the manuscript.

## Notes

### Competing Interest Statement

All authors are employees or contractors for VolitionRx. BA, JC, BPB, and TKK hold stock in VolitionRx.

### Summary of Updates

Update of Supplementary_Table file. There were some internal ID columns that were retained and unnecessary for external viewing.

